## Supplemental figures legend for "Nematode Small RNA Pathways in the Absence of piRNAs"

Supplemental Tables and Figures

**Tables:**

**Table S1. *Ascaris* small RNA pathway proteins and RNA expression**
**Table S2. *Ascaris* small RNA INPUT and IP libraries**
**Table S3. *Ascaris* small RNAs targeting repeats**
**Table S4. *Ascaris* small RNAs targeting mRNAs**

**Figures:**

**Figure S1. AsCSR-1 RNA expression and H3K9me3 marks.** A genome browser view on Chr17 shows the AsCSR-1 gene (in between 2.9 - 2.95 Mb), RNA-seq and H3K9me3 ChIP-seq from selected developmental stages. Note the high expression of AsCSR-1 in early embryos (1-4 cells) and the shutdown of its expression in 32-64 cells with heavy H3K9me3 marks.

**Figure S2. *Ascaris* Argonaute antibodies. A.** Western blots illustrating the specificity of antibodies and the relative amount of Argonaute proteins in the total cell lysate (T), cytoplasmic (C), or nuclear (N) fraction. The size of the Argonaute proteins is at ~ 100 kDa. An H3 histone variant (CENP-A) is used as a control for the cell fractionation. **B.** Argonautes IP small RNAs and Northern blots. Illustrated are small RNAs that co-IP with selected Argonautes antibodies on gels or from Northern blots. Note the high specificity of AsALG-1 IP for miRNAs and AsWAGO-1 for 22G-RNAs. CBP80 is a negative control that does not bind to any small RNAs. miR-100 sequence: AACCCGTAGATCCGAACTTGTGTT; Northern probe: CACAAGTTCGGATCTACGG. 22G-49 sequence: GTTAACGTAGAGCTCGCCAGAG; Northern probe: TGGCGAGCTCTACGTTAAC.

**Figure S3. *Ascaris* testes 5' all-phosphate vs. 5'-monophosphate small RNA libraries.** Comparison of small RNA library preparation methods (see methods) that capture 5' all-phosphate vs. 5'-monophosphate in broad regions of the testis. Region 1 corresponds to the mitotic region, transition zone, and early pachytene, Region 2 to late pachytene and meiosis, and Region 3 to late meiosis and mature spermatids. Note the enrichment of 5'-monophosphate small RNAs on right, i.e. miRNAs in Region 1 and 26G-RNAs in Region 2.

**Figure S4. *Ascaris* chromosome small RNAs.** Distribution of small RNAs associated with Argonautes on *Ascaris* chromosomes. Note the relative even distribution of AsCSR-1 associated small RNAs (targeting mRNAs) and the more biased distribution of AsWAGO-1 associated small RNAs (targeting repeats) in the genomic regions that will be eliminated during programmed DNA elimination. These eliminated regions have higher levels of repetitive sequences. Also note that the sex chromosomes are largely silenced in the male germline, thus there is a lack of siRNAs associated with AsCSR-1 on these chromosomes in the testis.

**Figure S5. Small RNA distribution on mRNAs.** Meta-analysis of siRNA distribution across mRNAs (in 100 bins on x-axis). The relative amount of siRNAs is shown on y-axis. Note the biased targeting at the 5'-end of mRNAs at late pachytene and meiosis for many Argonautes associated small RNAs, suggesting a possible concerted effort to repress the mRNAs for their clearance. Blue = antisense and Red = sense RNAs.

**Figure S6. Small RNA size distributions in INPUT and IP libraries.** The size distribution and frequency of small RNAs associated with INPUT or specific Argonaute IPs. Small RNAs (18-30 nt) from input and Argonaute IP were plotted for the whole sequencing library (All-siRNAs) or different types of siRNAs. Small RNAs starting with A, C, G, and U of different sizes (x-axis) were plotted against their read frequencies (y-axis; raw reads in millions). For each library, the y-axis scale is set the same as All-siRNAs to provide a direct comparison between different types of siRNAs within the same library.

**Figure S7. Comparisons of siRNA levels and their target RNA expression.** Pairwise comparisons of siRNAs associated with different argonautes and their targets. **A.** Repetitive sequences targeted in the testis. Note the similarity between AsWAGO-1, AsWAGO-2 and AsNRDE-3 and the overall repression of expression of their repeat targets. **B.** mRNA targets from testis in late pachytene and meiosis (M6). The red indicates mRNAs that are specifically expressed in male meiosis. Note the differential targeting between AsCSR-1 and AsALG-4 and the overlap between AsNRDE-3 and AsCSR-1. The targeted mRNAs show complex RNA expression levels, suggesting a potential mix of licensing, tuning and repressive functions.

**Figure S8. Argonaute associated small RNAs targeting Argonaute mRNAs.** Genome browser tracks showing four Argonautes that are targeted by siRNAs associated with these Argonautes throughout developmental stages. This illustrated auto-regulation of the expression of Argonautes by their associated small RNAs. For example, AsALG-4 is meiosis-specific and its expression in M5-M7 leads to the generation of 26G-RNAs that also target and repress AsALG-4 mRNA in M6-M7.
