## Supplemental figures for "Nematode Small RNA Pathways in the Absence of piRNAs"

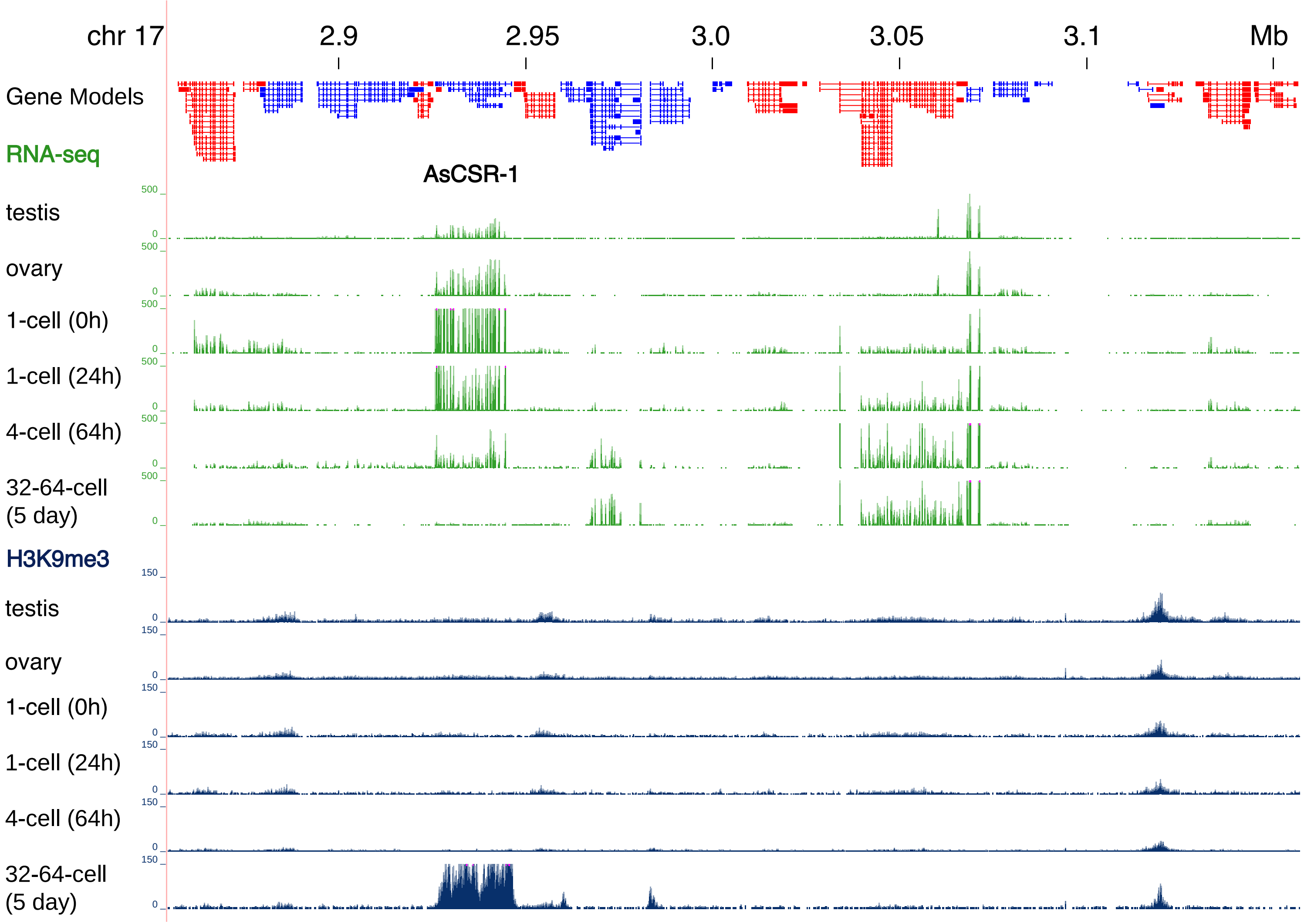

chr 17
2.9
2.95
3.0
3.05
3.1
Mb
Gene Models
RNA-seq
AsCSR-1
testis
ovary
1-cell (0h)
1-cell (24h)
4-cell (64h)
32-64-cell (5 day)
H3K9me3
testis
ovary
1-cell (0h)
1-cell (24h)
4-cell (64h)
32-64-cell (5 day)
500
0
150

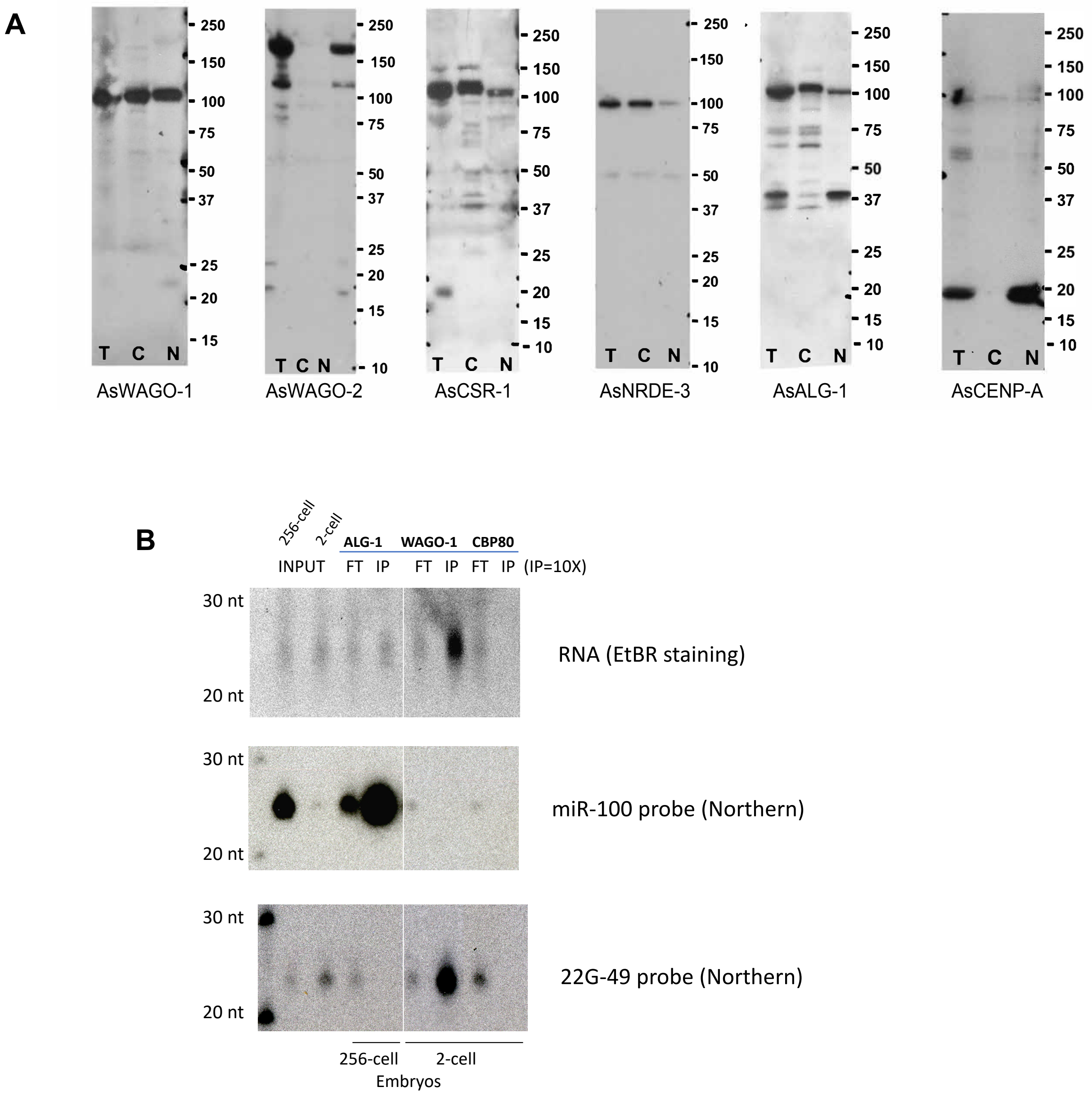
A
250
150
100
75
50
37
25
20
15
10
T C N
AsWAGO-1
AsWAGO-2
AsCSR-1
AsNRDE-3
AsALG-1
AsCENP-A
B
256-cell
2-cell
ALG-1
WAGO-1
CBP80
INPUT
FT IP FT IP FT IP (IP=10X)
30 nt
20 nt
RNA (EtBR staining)
miR-100 probe (Northern)
22G-49 probe (Northern)
256-cell
2-cell
Embryos

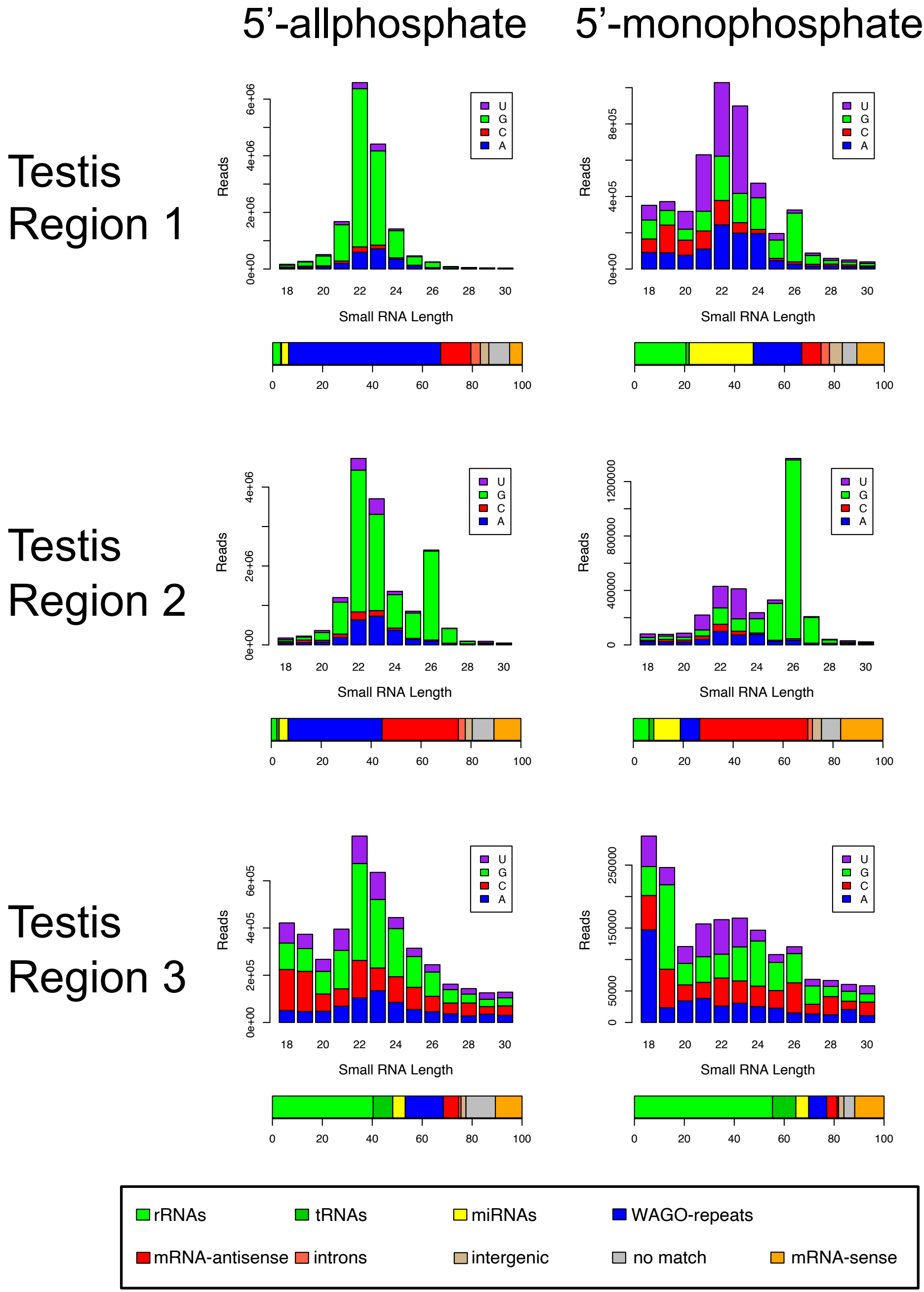

5'-allphosphate
5'-monophosphate
Testis Region 1
Testis Region 2
Testis Region 3
Reads
Small RNA Length
U
G
C
A
0e+00
2e+06
4e+06
6e+06
8e+05
4e+05
400000
800000
1200000
2e+05
6e+05
0
50000
150000
250000
18
20
22
24
26
28
30
0
20
40
60
80
100
rRNAs
tRNAs
miRNAs
WAGO-repeats
mRNA-antisense
introns
intergenic
no match
mRNA-sense

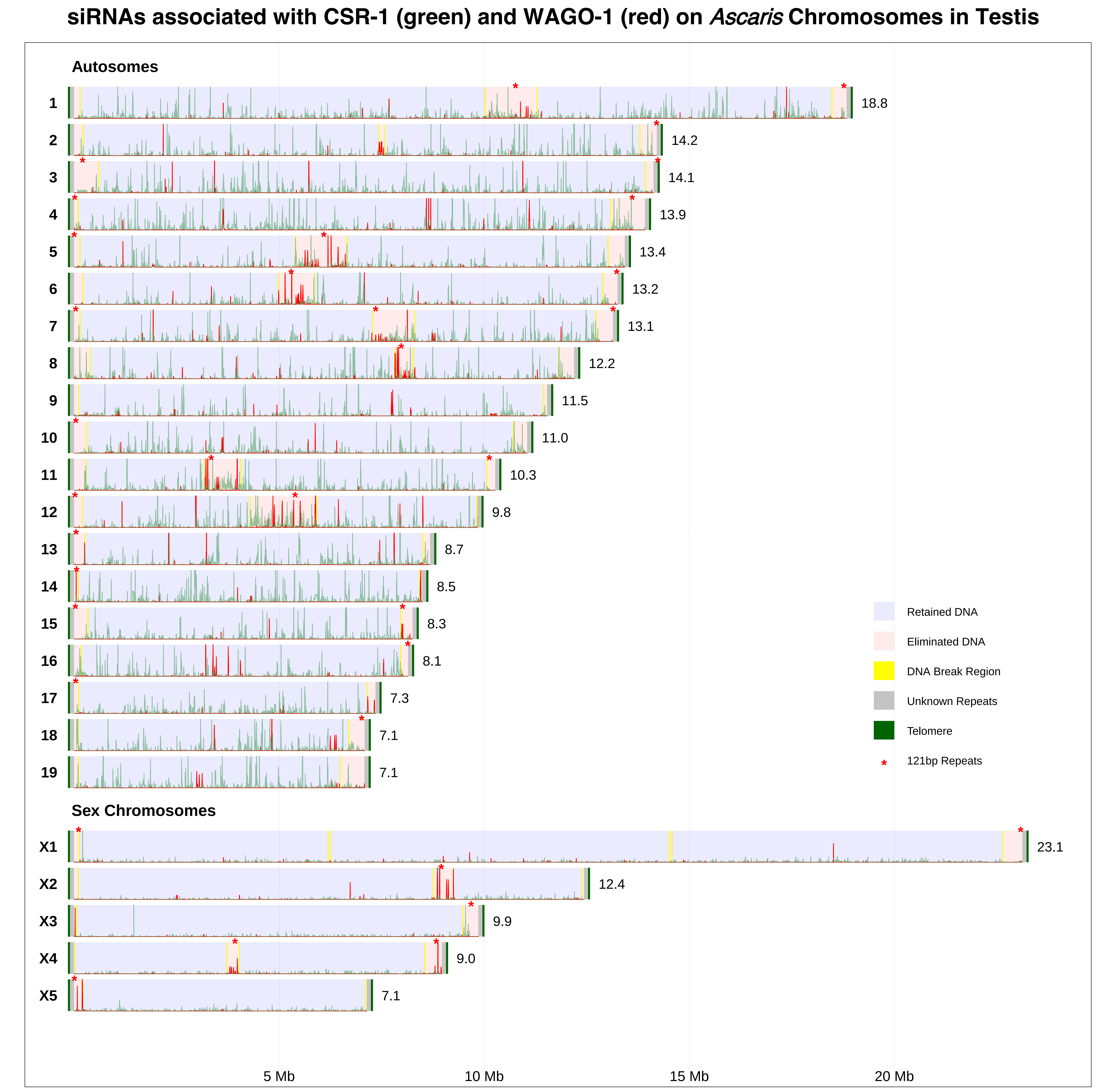

siRNAs associated with CSR-1 (green) and WAGO-1 (red) on *Ascaris* Chromosomes in Testis
Autosomes
1
18.8
2
14.2
3
14.1
4
13.9
5
13.4
6
13.2
7
13.1
8
12.2
9
11.5
10
11.0
11
10.3
12
9.8
13
8.7
14
8.5
15
8.3
16
8.1
17
7.3
18
7.1
19
7.1
Sex Chromosomes
X1
23.1
X2
12.4
X3
9.9
X4
9.0
X5
7.1
Retained DNA
Eliminated DNA
DNA Break Region
Unknown Repeats
Telomere
121bp Repeats
5 Mb
10 Mb
15 Mb
20 Mb

siRNAs associated with CSR-1 (green) and WAGO-1 (red) on *Ascaris* Chromosomes in Ovary

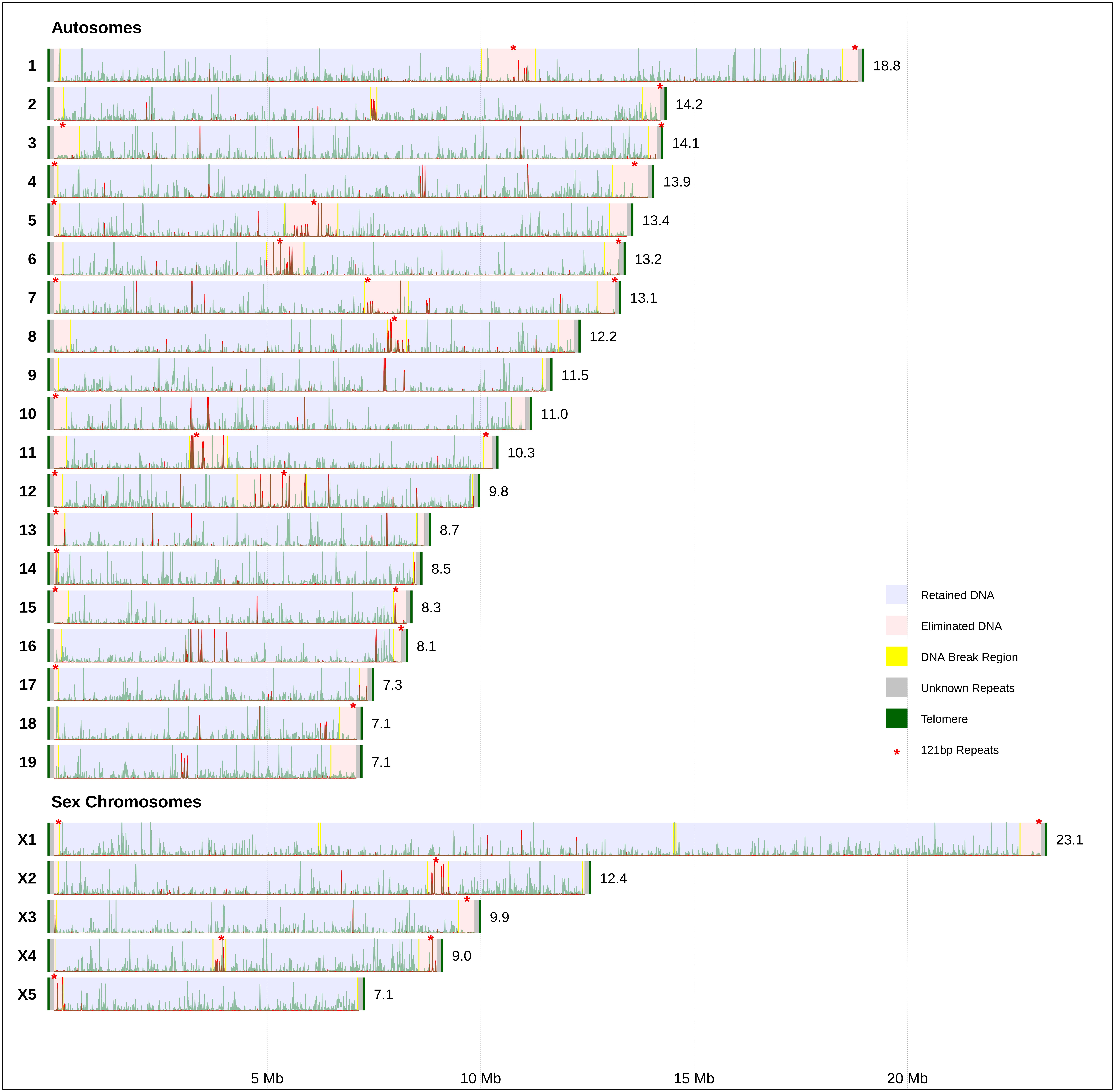

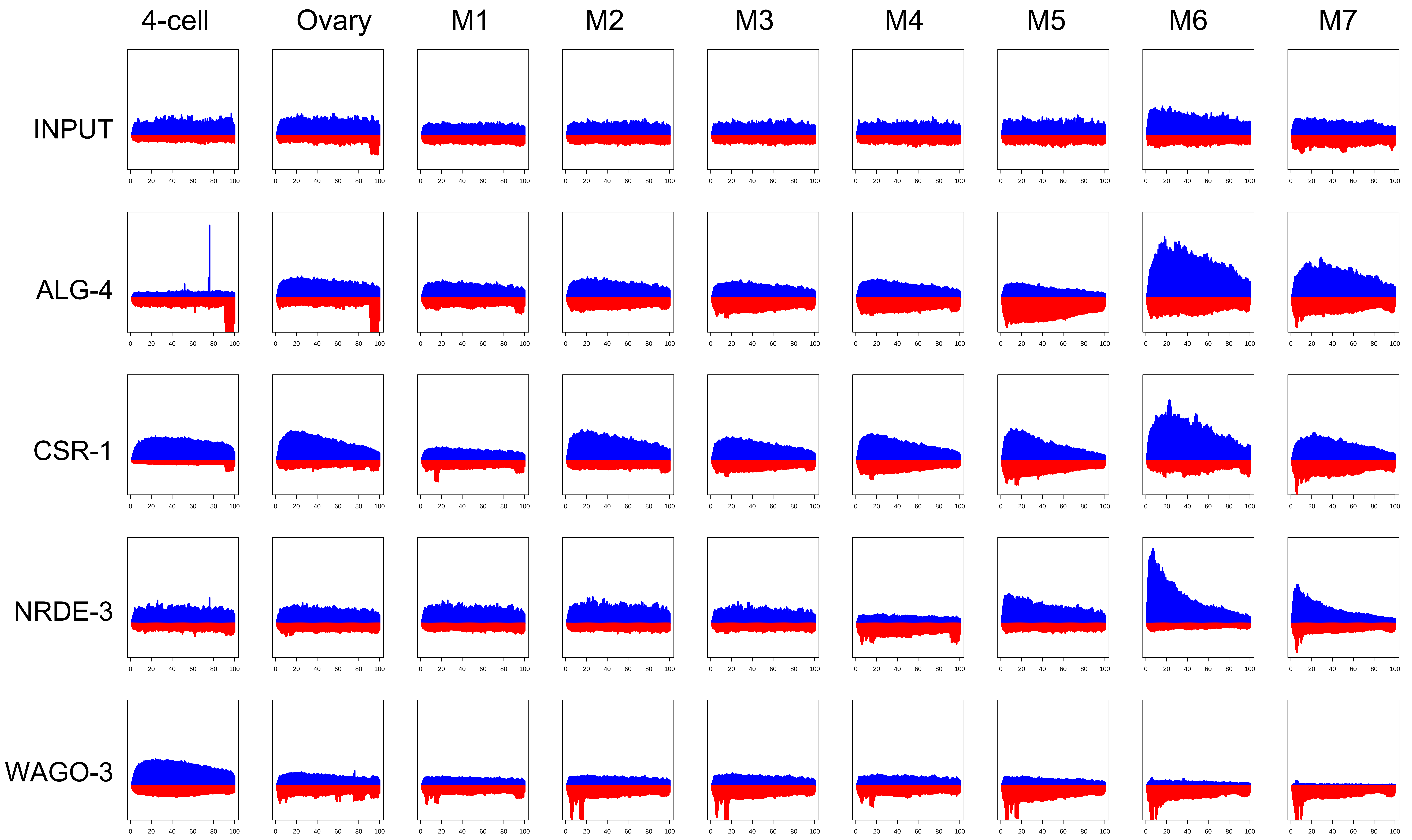
4-cell
Ovary
M1
M2
M3
M4
M5
M6
M7
INPUT
ALG-4
CSR-1
NRDE-3
WAGO-3
0 20 40 60 80 100

Fig. S6a

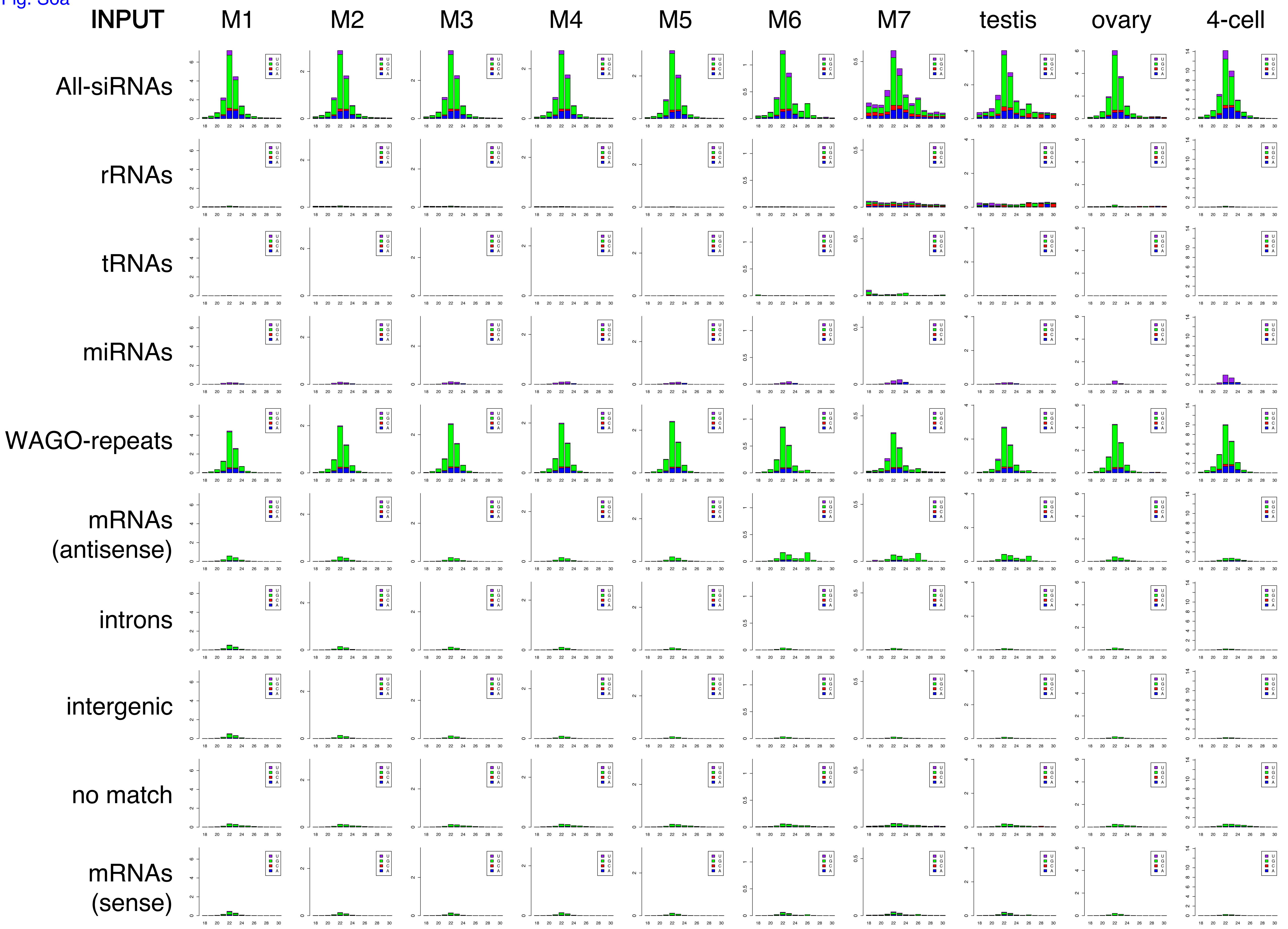
INPUT
M1
M2
M3
M4
M5
M6
M7
testis
ovary
4-cell
All-siRNAs
rRNAs
tRNAs
miRNAs
WAGO-repeats
mRNAs (antisense)
introns
intergenic
no match
mRNAs (sense)

Fig. S6b

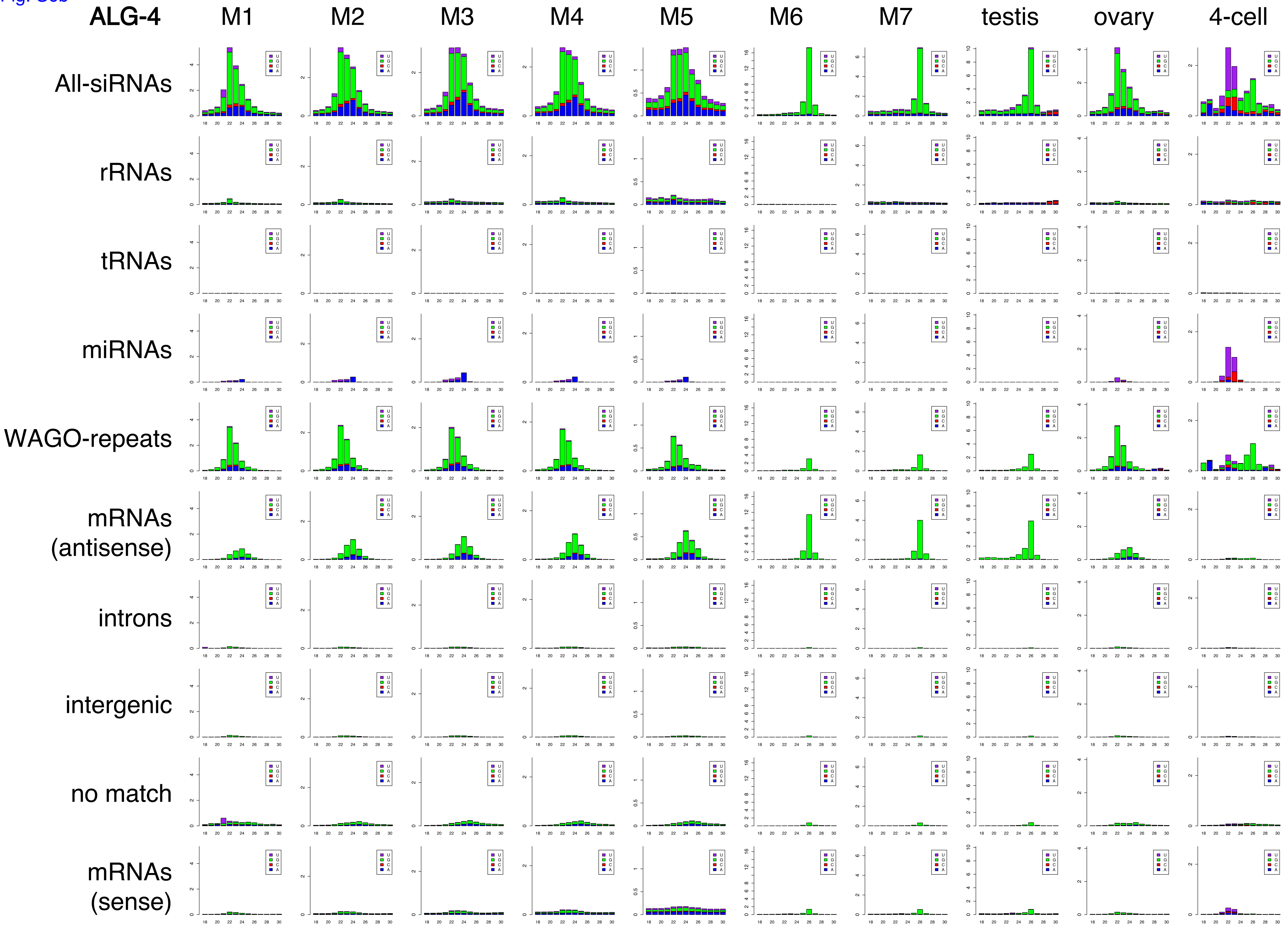
ALG-4
M1
M2
M3
M4
M5
M6
M7
testis
ovary
4-cell
All-siRNAs
rRNAs
tRNAs
miRNAs
WAGO-repeats
mRNAs (antisense)
introns
intergenic
no match
mRNAs (sense)
U
G
C
A

Fig. S6c

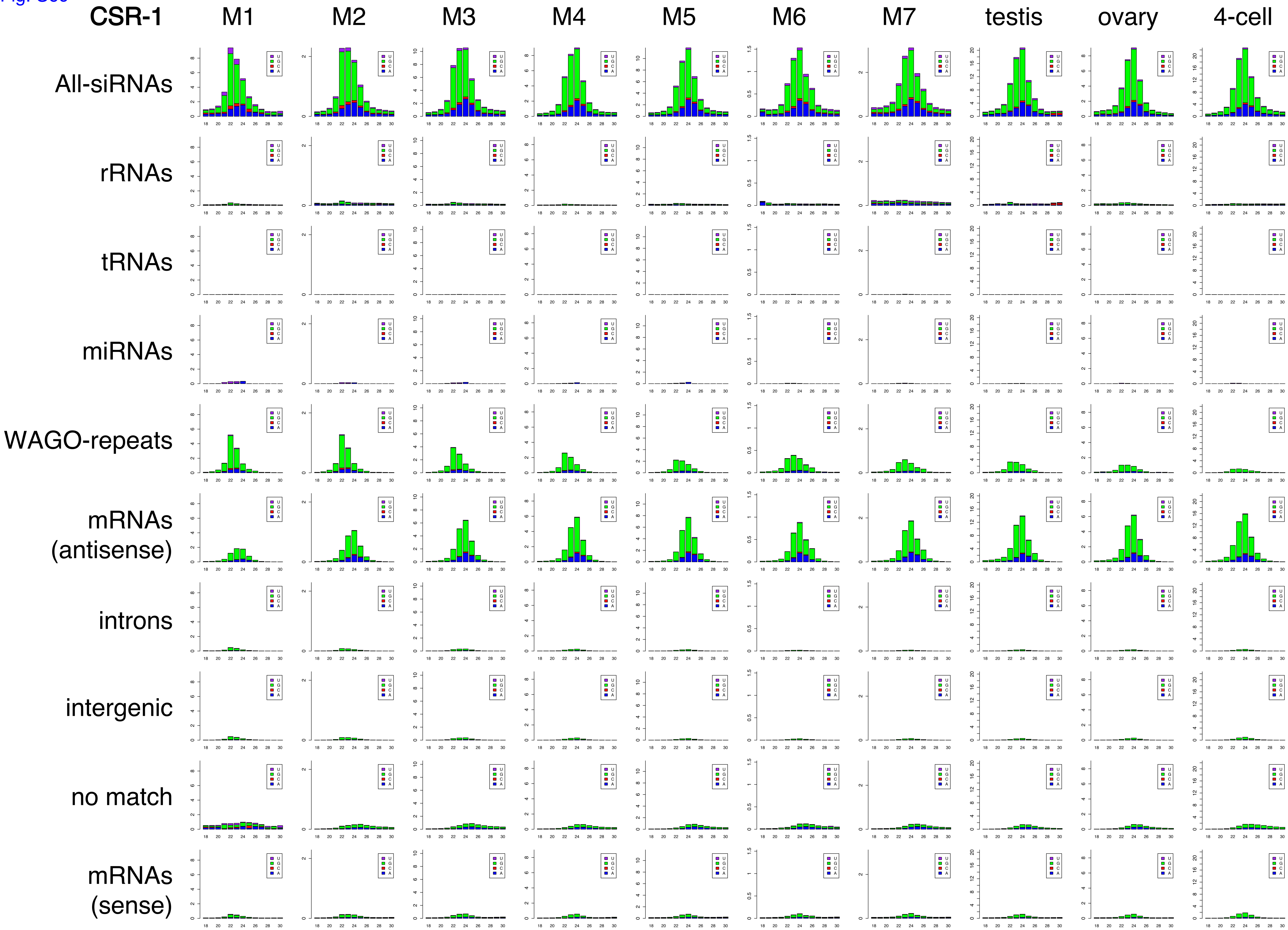
CSR-1
M1
M2
M3
M4
M5
M6
M7
testis
ovary
4-cell
All-siRNAs
rRNAs
tRNAs
miRNAs
WAGO-repeats
mRNAs (antisense)
introns
intergenic
no match
mRNAs (sense)
U
G
C
A

Fig. S6d

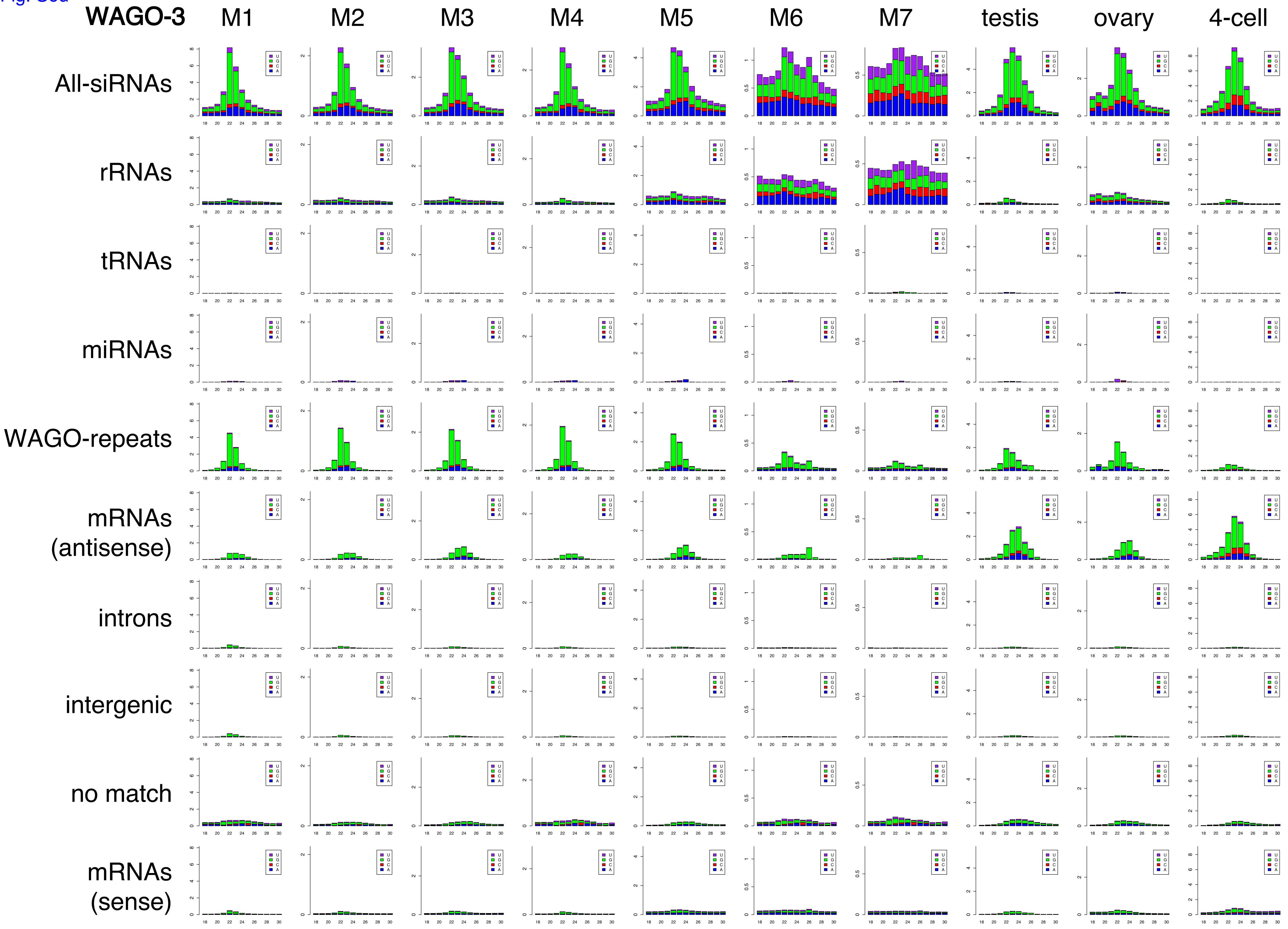
WAGO-3
M1
M2
M3
M4
M5
M6
M7
testis
ovary
4-cell
All-siRNAs
rRNAs
tRNAs
miRNAs
WAGO-repeats
mRNAs (antisense)
introns
intergenic
no match
mRNAs (sense)
U
G
C
A

Fig. S6e

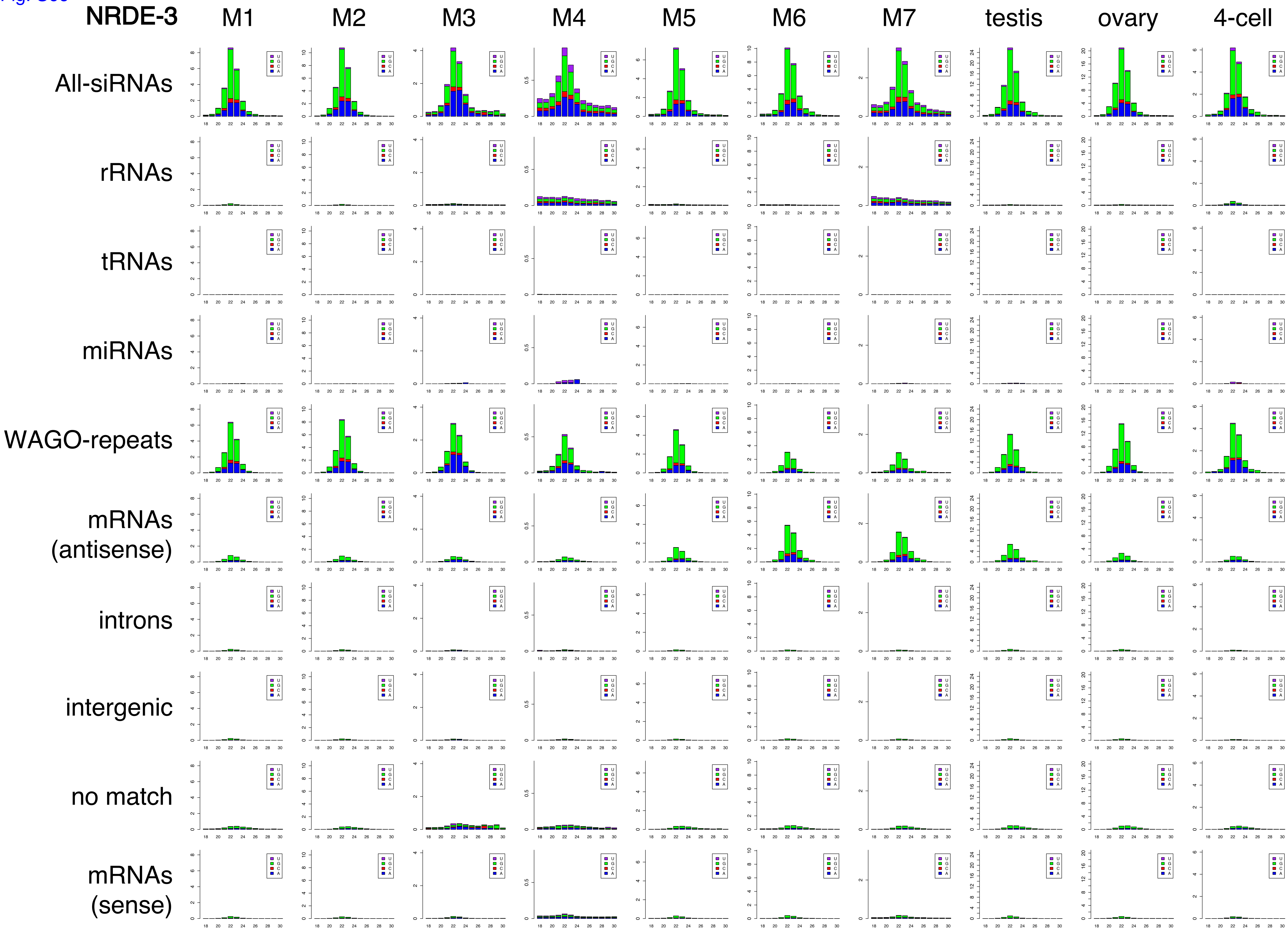
NRDE-3
M1
M2
M3
M4
M5
M6
M7
testis
ovary
4-cell
All-siRNAs
rRNAs
tRNAs
miRNAs
WAGO-repeats
mRNAs (antisense)
introns
intergenic
no match
mRNAs (sense)
U
G
C
A

Fig. S6f

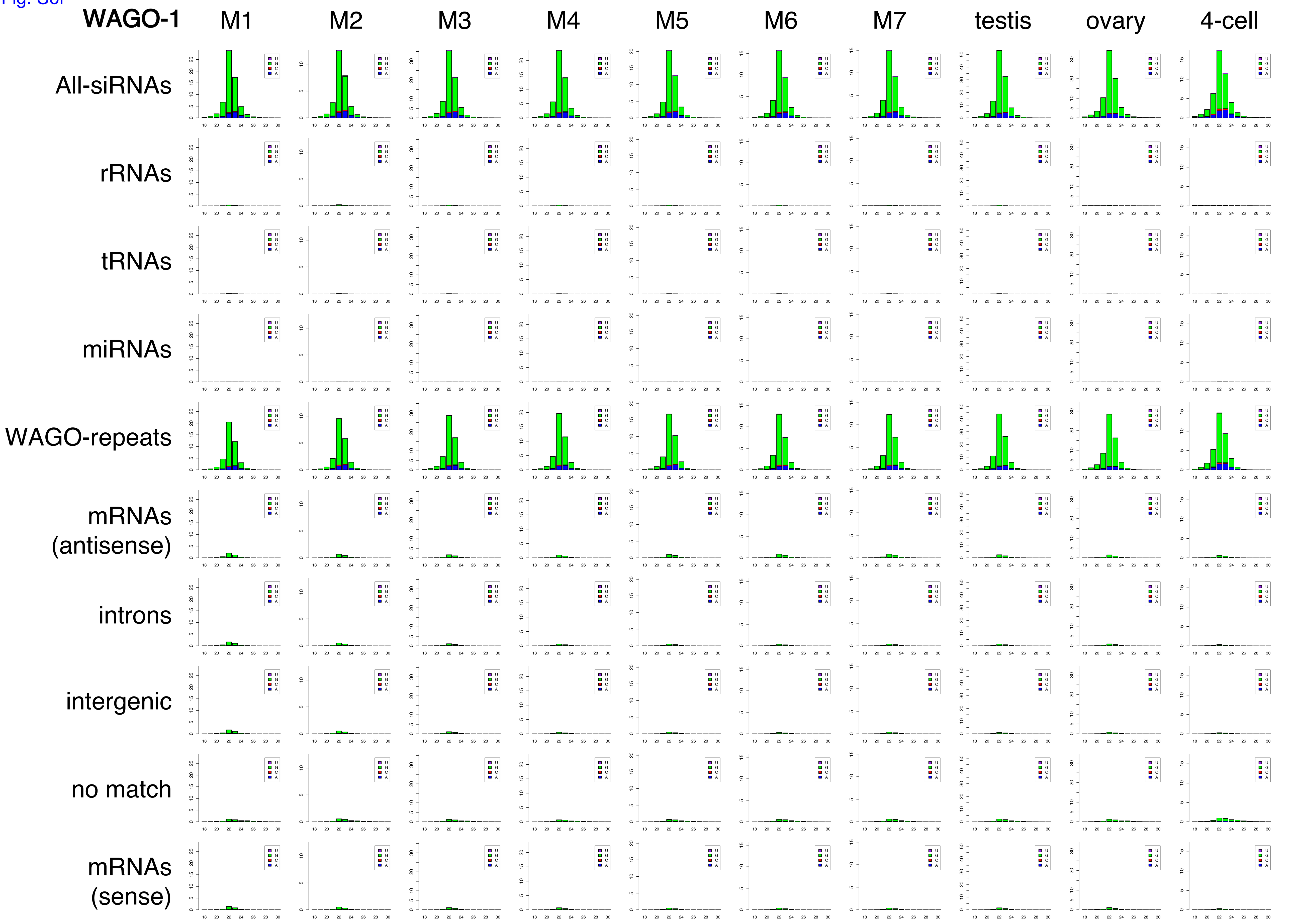
WAGO-1
M1
M2
M3
M4
M5
M6
M7
testis
ovary
4-cell
All-siRNAs
rRNAs
tRNAs
miRNAs
WAGO-repeats
mRNAs (antisense)
introns
intergenic
no match
mRNAs (sense)

Fig. S6g

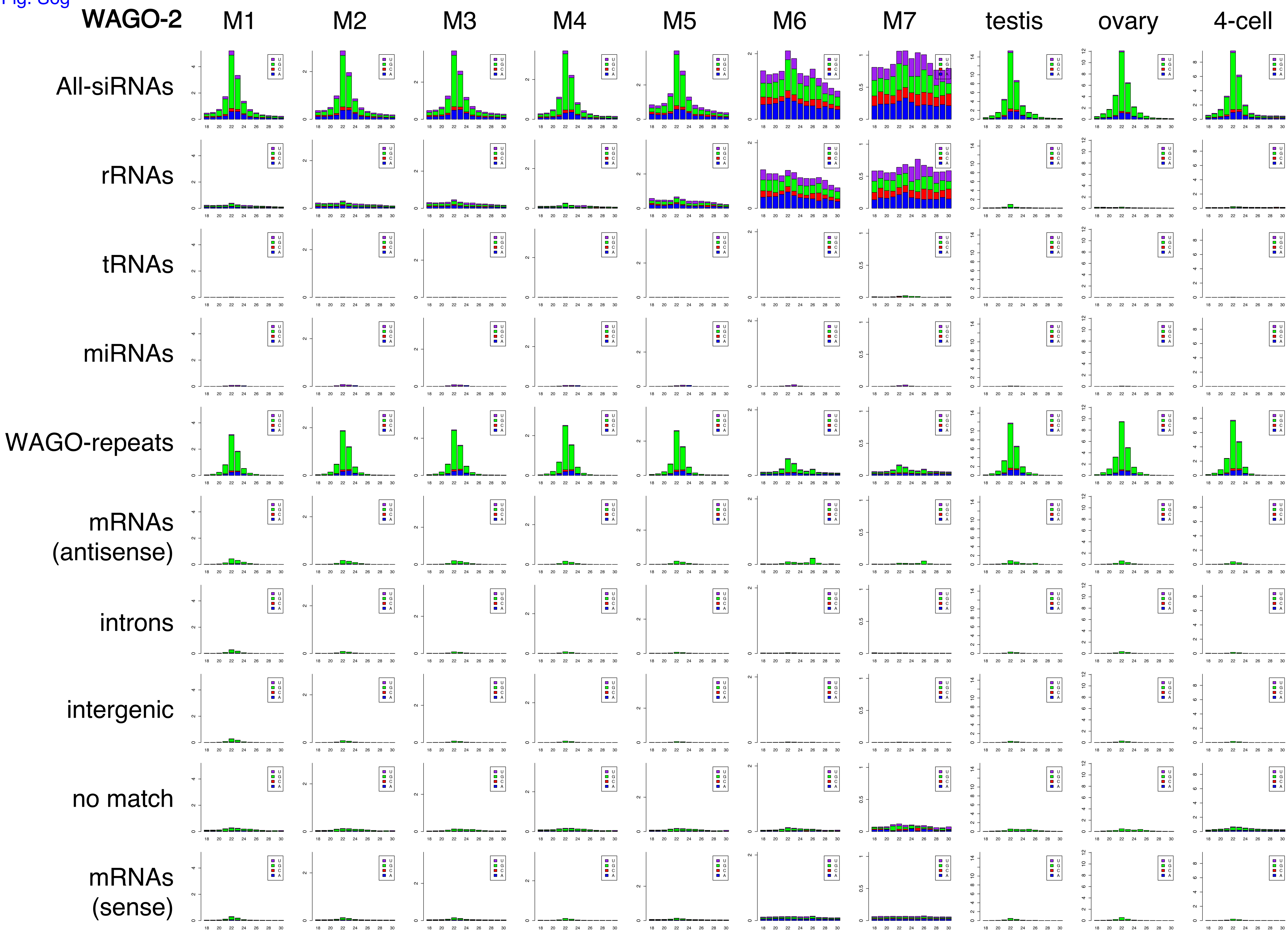
WAGO-2
M1
M2
M3
M4
M5
M6
M7
testis
ovary
4-cell
All-siRNAs
rRNAs
tRNAs
miRNAs
WAGO-repeats
mRNAs (antisense)
introns
intergenic
no match
mRNAs (sense)
U
G
C
A

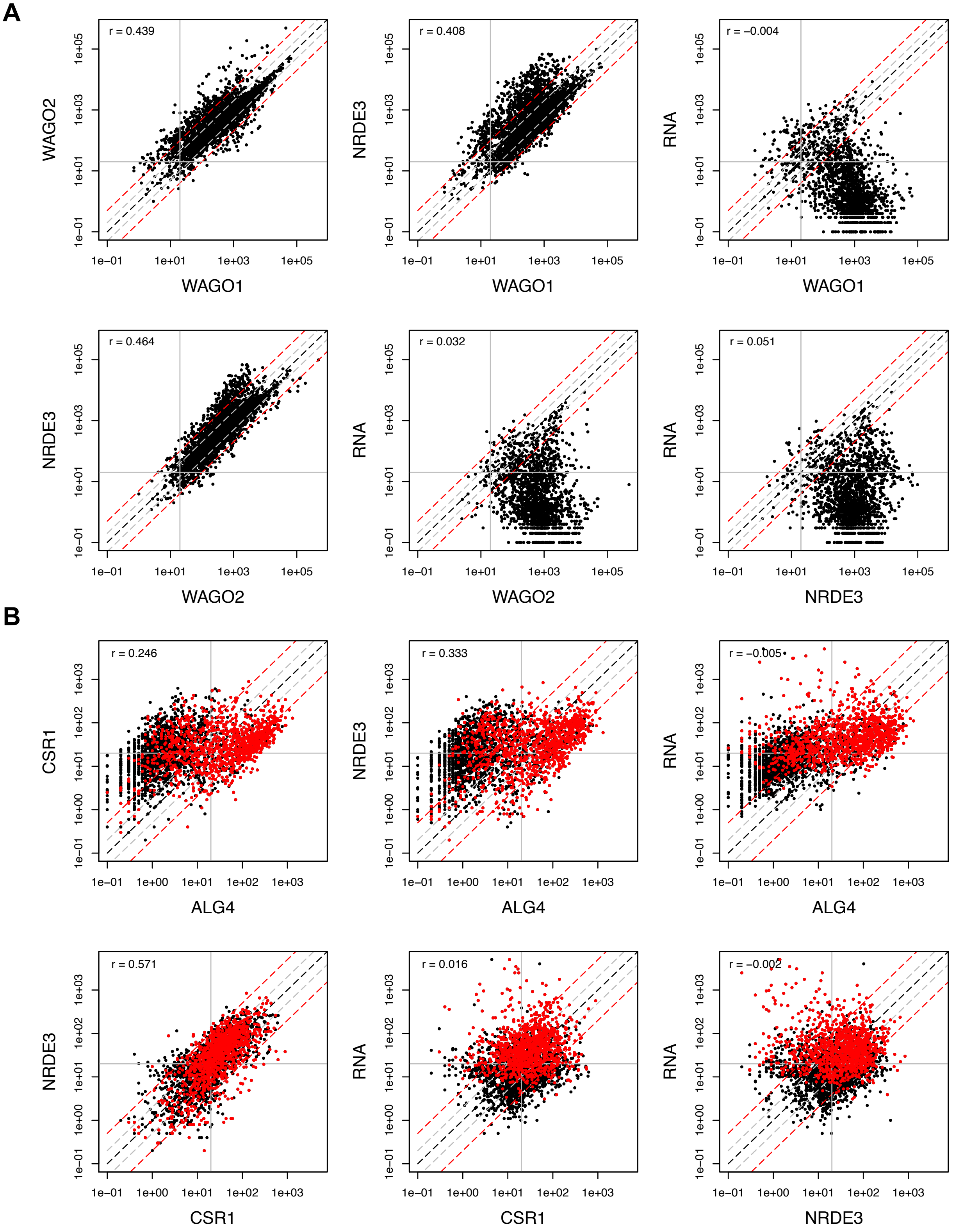
A
r = 0.439
WAGO2
WAGO1
r = 0.408
NRDE3
WAGO1
r = −0.004
RNA
WAGO1
r = 0.464
NRDE3
WAGO2
r = 0.032
RNA
WAGO2
r = 0.051
RNA
NRDE3
1e−01
1e+01
1e+03
1e+05
B
r = 0.246
CSR1
ALG4
r = 0.333
NRDE3
ALG4
r = −0.005
RNA
ALG4
r = 0.571
NRDE3
CSR1
r = 0.016
RNA
CSR1
r = −0.002
RNA
NRDE3
1e−01
1e+00
1e+01
1e+02
1e+03

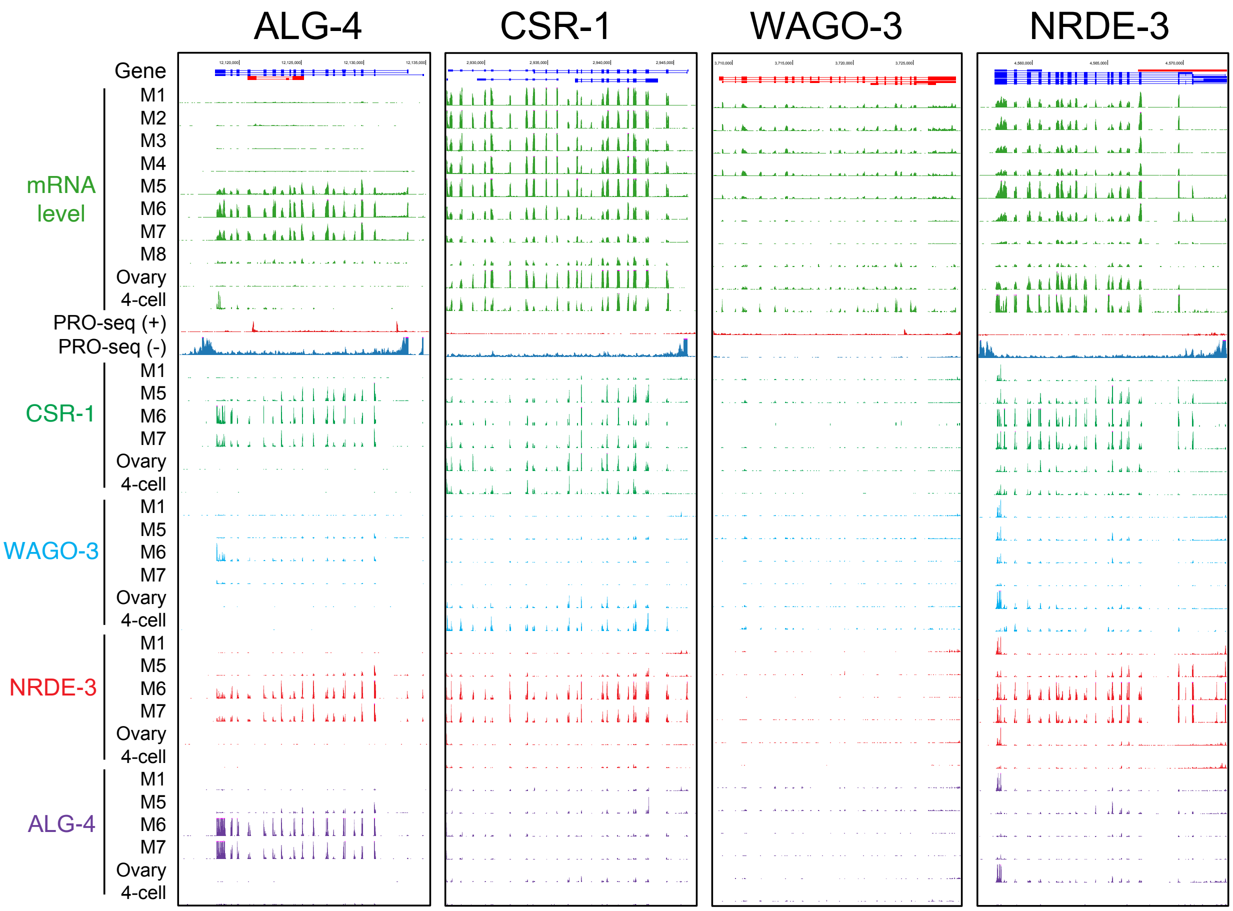

ALG-4
CSR-1
WAGO-3
NRDE-3
Gene
M1
M2
M3
M4
M5
M6
M7
M8
Ovary
4-cell
mRNA level
PRO-seq (+)
PRO-seq (-)
CSR-1
M1
M5
M6
M7
Ovary
4-cell
WAGO-3
M1
M5
M6
M7
Ovary
4-cell
NRDE-3
M1
M5
M6
M7
Ovary
4-cell
ALG-4
M1
M5
M6
M7
Ovary
4-cell
